# Bacterial community characterization by deep learning aided image analysis in soil chips

**DOI:** 10.1101/2023.11.13.566656

**Authors:** Hanbang Zou, Alexandros Sopasakis, François Maillard, Erik Karlsson, Julia Duljas, Simon Silwer, Pelle Ohlsson, Edith C. Hammer

## Abstract

Soil microbes play an important role in governing global processes such as carbon cycling, but it is challenging to study them embedded in their natural environment and at the single cell level due to the opaque nature of the soil. Nonetheless, progress has been achieved in recent years towards visualizing microbial activities and organo-mineral interaction at the pore scale, especially thanks to the development of microfluidic ‘soil chips’ creating transparent soil model habitats. Image-based analyses come with new challenges as manual counting of bacteria in thousands of digital images taken from the soil chips is excessively time-consuming, while simple thresholding cannot be applied due to the background of soil minerals and debris. Here, we adopt the well-developed deep learning algorithm Mask-RCNN to quantitatively analyse the bacterial communities in soil samples from different locations in the world. This work demonstrates analysis of bacterial abundance from three contrasting locations (Greenland, Sweden and Kenya) using deep learning in microfluidic soil chips in order to characterize population and community dynamics. We additionally quantified cell- and colony morphology including cell size, shape and the cell aggregation level via calculation of the distance to the nearest neighbor. This approach allows for the first time an automated visual investigation of soil bacterial communities, and a crude biodiversity measure based on phenotypic cell morphology, which could become a valuable complement to molecular studies.

## 1 INTRODUCTION

Direct investigation of soil processes involving microbes, trophic interactions and organic matter cycling at the microscale has been a major challenge due to the opacity of soil and the exceedingly complex pore spaces. Important processes such as carbon cycling are studied intensively but need to rely on bulk- and black-box approaches, and even though we know that microbes play a major role in driving the carbon cycle we commonly cannot study them at the mechanistic level at the scale of their cells Hill et al. (2000); Chatterjee et al. (2009). Soil chips are microfabricated microfluidic devices typically made of glass and polydimethylsiloxane (PDMS), designed to mimic the natural soil pore network to address the spatial structure that commonly is missing in laboratory experiments, and allow real-time visualization and characterization of microbial activity via non-destructive and label free methods such as optical microscopy, Raman scattering and synchrotron radiation based X-ray microspectroscopy at the micro and nano scale Pucetaite et al. (2021); Aleklett et al. (2018). They have so far been used to study soil microbial questions, including investigating microbial communities from real soil Nichols et al. (2010); Mafla-Endara et al. (2021) and studying the effect of soil structure on microbial activities and carbon cycling Arellano-Caicedo et al. (2021); Aleklett et al. (2021); Soufan et al. (2018); Deng et al. (2015); Ceriotti et al. (2022); Aufrecht et al. (2019).

Due to the transparent nature of PDMS and glass, bacterial and fungal growth, behaviour and substrate degradation can be directly visualized through optical microscopy at real time. Quantification and characterization of microorganisms can be easily achieved using bacterial or fungal strains and substrates with fluorescence labels. Natural soil communities can also be used as inoculum and investigated within soil chips, allowing for the observation and tracking of live interactions and growth using non-destructive and label-free methods which is bright-field imaging via an optical microscope. The images acquired from natural communities in soil chips normally consist of complex microstructures and different microorganisms plus soil minerals and debris, which makes it an intricate task to differentiate the microorganisms from the complex background. There are existing software and scripts for bacteria counting but they mostly work for clean cultures Grishagin (2015); Selinummi et al. (2005); Ferrari et al. (2017) and the performance on soilchip tasks is not satisfying. So far, counting tasks have been carried out manually, but this may lead to high variance and bias from human errors Chien et al. (2007) and is laborious. Automated counting can be more efficient and reproducible. It requires the segmentation of the objects followed by classification. Interestingly, soil microbes in microscopy images may not have a clear detectable boundary for thresholding but they typically have a discernible morphology from the background and microstructure of the chips, requiring deep learning approaches. Deep learning object recognition is proven effective and efficient for identifying and extracting features thanks to the development of the GPUs computing power and deep convolution neural networks (CNN), a biologically inspired deep learning algorithm Jiao et al. (2019). Bacterial population sizes have been quantified at a satisfying accuracy of CNN-based classification architecture, although the images were taken from culture plates Ferrari et al. (2015, 2017); Tamiev et al. (2020); Naets et al. (2021); Spahn et al. (2021). Zhang et al. has rigorously organized and reviewed existing image-based bacteria counting from manual to AI and extrapolated that computer vision based microorganism counting methods will replace traditional method as it combines both segmentation and classification along with high accuracy Zhang et al. (2021).

In this study, we trained a model using the Matterport implementation of Mask RCNN for bacterial recognition on the images from our experimental data Abdulla (2017). Mask R-CNN He et al. (2017) is an extension from Faster R-CNN Girshick (2015) for instance segmentation tasks with advantages of high localization and objects recognition accuracy. It has been extensively applied for disease detection and extraction of various other features Johnson (2018); Khan et al. (2021); Zhang et al. (2020); Chiao et al. (2019); Couteaux et al. (2019). We utilized the model to analyze data acquired from three contrasting locations, including Lund, Sweden, Nyabeda, Western Kenya, and Disko research station, Greenland. These locations exhibit a significant variance in environmental conditions, ranging from extreme climates and low nutrient availability in a high arctic biocrust to a fertile temperate grassland and a tropical agricultural system, using different sample preparations and microscopy techniques. Our study aimed to explore the potential of deep learning algorithms in bacterial image recognition of environmental samples using these datasets and provide guidelines for dataset preparation and training. With the increasing user-friendliness of deep learning for biology groups without much AI experience, we found that image annotation equivalent to a few weeks’ work can provide satisfactory information on bacterial populations within the chips. This method is expected to be useful for environmental microbiological research and analysis using imaging methods, particularly on microfluidic platforms, where the combination of controlled experimental manipulations and monitoring of processes with high environmental realism, including chemo-physical conditions and biotic interactions, presents significant potential.

## 2 EXPERIMENTAL METHODS

### 2.1 Soil sample acquisition

The deep learning model was tested with image data of varying quality, obtained from soil chips inoculated from three contrasting soil/soil-like inocula: a biocrust from Disko research station, Greenland; a moraine clay till from Lund, southern SwedenDaniel et al. (2000); and a Lixic Ferrasol from Nyabeda, Siaya County, Western KenyaOmuto (2013). All the image acquisition was carried out via optical microscopy either in lab or *in situ* without staining and labeling as a non-destructive and label free approach is required for minimal intervention. The images of samples from Sweden and Greenland were taken in the lab whereas the images of samples from Kenya were taken *in situ*. The samples from Greenland were collected during the summer of 2021 from biological soil crusts and were incubated as intact crust parts (1 x 5 *cm*^2^ dimensions, 1 cm height) on liquid malt-medium filled chips first in a room temperature condition for a week, and then moved to phytotron (adaptis by CONVIRON) at 5^*°*^C and 80 % humidity for storage.

The samples from Sweden were collected during the summer of 2021 at a depth of 10 cm from a lawn outside the Ecology Building at Lund University. The soil samples were sieved through a 2 mm mesh strainer and stored in a plastic zip-lock bag at 4^*°*^C. 10 grams of soil sample were inoculated onto the malt-medium filled microfluidic device opening to initiate the colonization of microbial communities into the environment, incubated at room temperature in dark. The chips were monitored and images of bacterial populations were taken over 88 days for the Greenland samples and over 28 days for the Sweden samples, both acquired using a Nikon Ti2-E inverted light microscope with a Nikon Qi2 camera in the lab.

In Kenya, the data was collected during fieldwork in spring 2022 from unearthed chips *in situ* placed into cropland fields planted with maize (Zea mays) as part of a maize–soybean crop rotation scheme, located in Nyabeda, Siaya County by the Lake Victoria basin Kätterer et al. (2019). The M9-medium filled soil chips were buried at a depth of 10-20 cm, and 10-20 cm away from the crop row for 49-56 days. Upon the retrieval of the chips from the soil, images were acquired on the spot with a portable inverted field microscope (EmMicroscope with x10 N/A 0.25 and x40 N/A 0.65 objectives). Photos were captured using a Samsung S20 FE 5G SM-G781B phone, with the following camera settings: 12.2MP, 3024×4032, f/1.8, 1/1936, 5.4mm, ISO40.

The three sampling locations exhibit noticeable differences in bacterial abundance and morphology, which can be attributed to various experimental conditions and strong differences in image quality. These data were utilized as a validation set to assess the efficacy of our deep learning recognition model in accurately capturing and quantifying such differences, thereby demonstrating its proof of concept. The sampling sites were thus chosen to provide large differences in environmental conditions to demonstrate the versatility of the approach; it should be noted that our objective did not encompass a comparison of the effects of different experimental conditions on bacterial morphology.

### 2.2 Microfluidic chips

The microfluidic soil chips serve as idealized micro-structured habitats, featuring a micro-scale network of predefined poly-dimethylsiloxane (PDMS) structures that replicate soil pore spaces through photolithography. The utilization of soil chips allows for direct observation of active microorganisms migrating from natural soil samples into the artificial structures, facilitating the investigation of diverse microbial community interactions, including inter-kingdom, food-web interactions, and feedbacks between microbes and pore space microstructures Mafla-Endara et al. (2021).

In an additional experiment conducted to demonstrate the soil chip’s efficacy (refer to the supplementary information on bacterial metabarcoding for microfluidic soil chips), we observed a broad diversity within bacterial communities in the microfluidic soil chips, representing 26 to 39% of the soil bacterial richness (Table S1). Furthermore, there is a substantial overlap between bacterial communities developing in the soil chips and their soil inoculum (Table S2). This highlights the effectiveness and utility of the soil chips as a powerful tool for exploring and experimenting with diverse soil bacterial communities and their interactions, both biotic with other microbial kingdoms and abiotic experimental factors such as microstructures or toxins.

Before soil inoculation, the chips were filled with a liquid nutrient medium that allowed for microbial migration from the soil prior to the inoculation step. The inherent nutrients present in the soil can then freely diffuse into the chip as it is filled with liquid and oxygenation was ensured via the gas permeable PDMS polymer. The soil chips contain an experimental section and a entry system near the inoculation reservoir featuring identical cylindrical pillars designed to prevent the PDMS lid from collapsing and to facilitate microbial migration into the specific experimental section. The microorganisms actively migrate into the chips with a potential travel distance of 5 cm. Within a day, first bacteria can often be observed in the distant parts of the chips due to their active movement. Additionally, water movement during drying-rewetting processes can also drag organisms and mineral particles into the chips Mafla-Endara et al. (2021). All microorganisms and mineral particles that can fit into spaces of 12*µm* height can potentially enter the chips. The entry system commonly measures a depth of 3 to 6 mm until the specific experimental section. The design details for the experimental chips used in Sweden and Kenya are elaborated in work by Aleklett et al. (2021), while the specifics for the chips utilized in Greenland can be found in the study by Arellano-Caicedo et al. (2023).

For the experiments in Greenland and Sweden, images were taken on four chip replicates at different internal chip locations over varying time periods. In contrast, the dataset from Kenya is a single-time measurement with five replicates in the experimental section. Only the images taken from the entry system were used for comparison between Greenland and Sweden, which applies to the Figure 2 to 5 due to the same structures and distance to the soil inoculum. All the available images including the ones taken in the specific experimental section from Greenland were used to obtain the morphology information in Figure 6.

### 2.3 Module training and evaluation

We adapted an instance segmentation algorithm based on Mask-RCNN. We started the training of the algorithm on one of the datasets (images of bacteria from the Swedish soil) and then extended its recognition ability to the other two ecosystems and acquisition approaches, which differed in image quality and variation of bacterial morphology. The objects of the images were categorized into five classes: ‘microfluidic structures’, ‘fungal hyphae’, ‘single bacteria’, ‘dividing bacteria’ (two attached bacteria) and ‘bacterial clusters’ (more than two bacteria). As a limitation of morphological analysis, it is important to note that microorganisms with similar shapes, such as bacteria, archaea and small yeast, cannot be differentiated based on morphology alone. Therefore, when referring to the classification of “bacteria,” it should be interpreted as a shape object resembling that of bacteria rather than a conclusive identification of bacterial species. Although yeast cells could be mistaken for bacteria due to their similar morphology, we believe this is mostly insignificant in our experimental system. Yeasts typically account for only 0 to 10% of total soil fungal communities Mašínová et al. (2017), and the amount of fungal colonies forming units is consistently two to three orders of magnitude lower than that of bacterial cells in soilsKacergius and Sivojiene (2023); Kwiatkowska et al. (2022); Sofo et al. (2020). For these reasons, we consider the potential skewing of our bacterial results by yeast contamination to be negligible. Archaea were confirmed to be included in the microbial communities inside the soil chips (Supplemental Information). The ratio of bacteria to archaea in soil chips under different conditions should be investigated in future studies. Images were cropped to 512*X*512 *px* as a balance between GPU computing power and batch size. Those images were then annotated using the Viavgg website based image annotation software Dutta et al. (2016); Dutta and Zisserman (2019). The training of the algorithm was carried out through the Alvis supercomputer facility (a Swedish National Infrastructure for Computing (SNIC) resource dedicated to artificial intelligence and machine learning research). Following the methodology suggested in Naets et al. (2021), we split the data acquired from the experiment using Swedish soil into 65 images for training and 16 images for validation. We started with transfer learning from already annotated daily common objects dataset (COCO) on only head layers with RestNet 50 as backbone model Lin et al. (2014). Here, transfer learning refers to fine-tuning of models already trained on different tasks and datasets that have plenty of annotations. Subsequently, all the layers were trained for 500 epochs followed by another 500 epochs training with image augmentations which include flip, rotations and affine transformation. We then trained the same dataset with ResNet-101 network with heavier augmentation for 500 epochs. It is worth noting that the selection of image augmentations is of crucial importance to improve training performance. Multiply, linear contrast, sharpen, emboss, flip in horizontal and vertical directions, rotations and affine transformation were used for heavy augmentation.

Annotating dataset images is one of the most tedious and time-consuming parts of the modelling process. The size of the dataset is important - a large dataset is needed for model success but it takes a lot more time to label. In order to prepare a larger dataset to generalize the model for the Greenland data, the model was partially trained to learn the data first and then the model itself was used to annotate new data before training it further. The new dataset annotated by the partially trained model contains 540 training images and 141 validation images. The self-annotation dataset was manually checked and edited by a human. As a result, a large dataset can be prepared within a day. Since the image quality taken from the portable microscope in Kenya was much lower compared to the other two, a small dataset that only contained images from Kenya was used to fine tune the model for this data. Images from Kenyan soils were thus not included into the overall training dataset for the analysis of Swedish and Greenland data.

For the purpose of evaluating the performance of the model, two standard metrics are used to evaluate the accuracy of the detection: mean average precision (mAP) and recall. Mean average precision is the average of the weighted mean of precision of each class, and recall is the percentage of correct detected objects out of all the correct targets. In other words, recall represents the capability of detecting all the all instances of the target class and mean average precision shows how much we can trust the identified result.

Twenty images not previously used for training or validation were randomly selected from each dataset to evaluate the performance of the model. Dust or air-bubbles can look like bacteria using a monochromatic camera. Although the structure class is added to avoid this, the model sometimes detects some defects as bacteria inside the PDMS structure. Therefore, objects detected within the PDMS structures were removed afterwards. The result metrics are for the target objects ‘bacteria’, ‘dividing bacteria’ and ‘bacteria clusters’. As shown in Table 1, all the test images from Sweden and Greenland have gained satisfactory precision and recall. By lowering the minimum detection confidence threshold, the model tends to identify a greater number of objects, which increases recall by including more true and false positives; however, this also typically results in a decrease in precision due to the increase in false positives, reflecting the inherent trade-off between precision and recall. As shown in Figure 1, the model performs consistently over images with various brightness, contrast and bacteria at different focal depths compared to human labeling and is thus fit to replace human counting.

**TABLE 1.** Benchmarks (in percentage) of our best model without post image processing. Minimum detection confidence scores 0.7 is used.

| Iou threshold=0.5 | Greenland | Sweden | Kenya |
| --- | --- | --- | --- |
| Average precision | 90% | 91% | 89% |
| Average recall | 94% | 95% | 96% |

**FIGURE 1.**
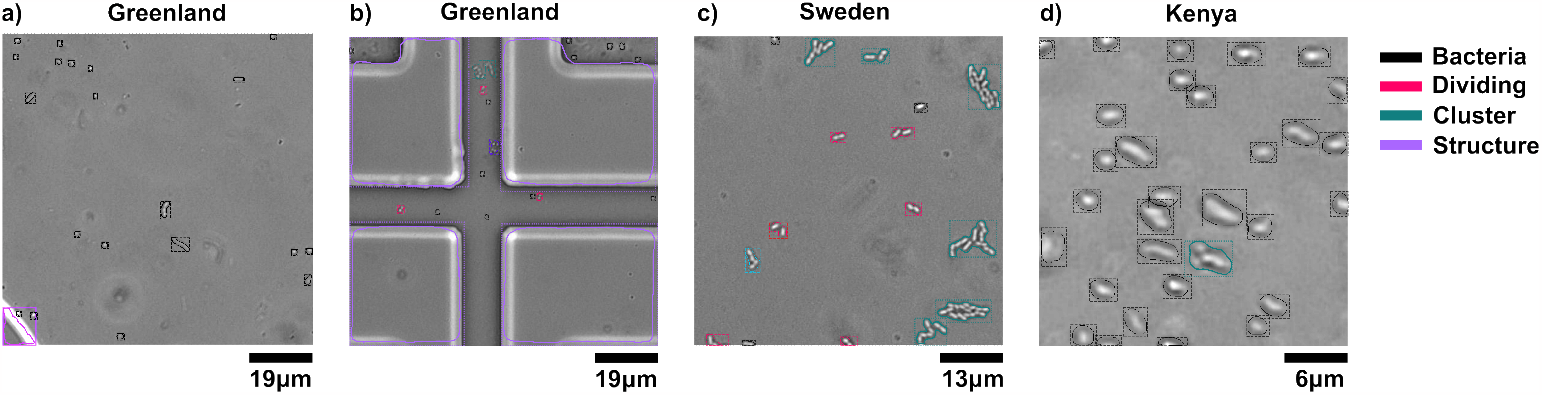
Example of labeled images taken from a-b) Greenland, c) Sweden and d) Kenya. The images are sized 512*X*512 *px* (thus the different levels of magnification). The color represents different classifications which are ‘bacteria’ (black), ‘dividing bacteria’ (pink), ‘bacterial cluster’ (green) and ‘structures’ (purple).

## 2.4 Data treatment

The number of bacteria for each replicate was estimated by averaging the number of the detected bacteria in each image (n=X). The average population density and standard error for each day were obtained by averaging the number of ‘bacteria’ over the chip replicates (n=4). The same method was used for calculating the average population density of the object classes ‘dividing’ and ‘cluster’.

To calculate the estimated total number of bacteria per unit area as displayed in Figure 3, the total number of detected bacteria was first calculated by adding the number of single bacteria and the estimated number of bacteria in the classes ‘dividing’ and ‘cluster’, and then divided by the available area (total area minus area occupied by microfluidic structures) in the space for each image. Each object classified as ‘dividing’ was considered as two bacteria and the area segmented as a ‘cluster’ was divided by the average size of bacteria from the detected image.

**FIGURE 2.**
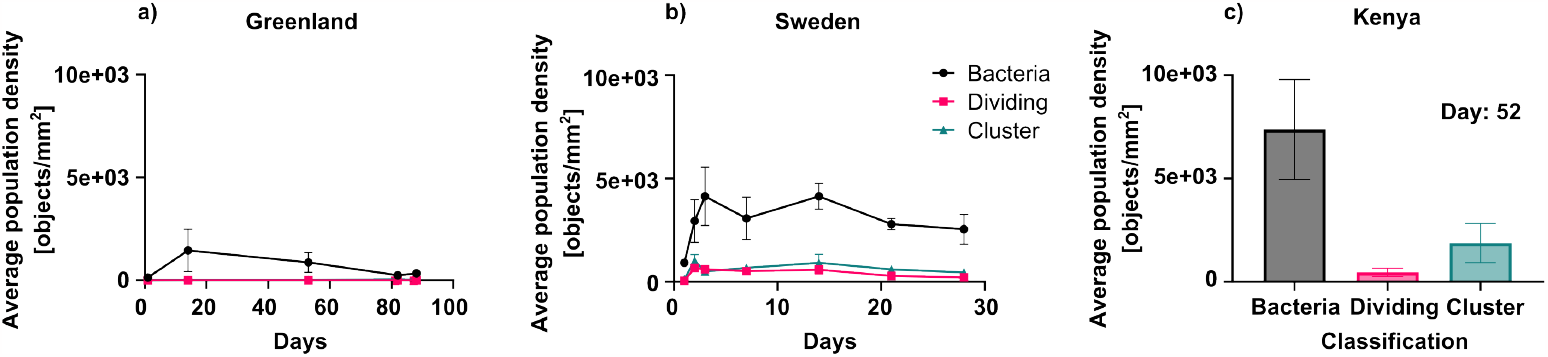
a,b) Average population density of detected objects as a function of time after inoculation in Greenland and Sweden for the three classifications, namely ‘bacteria’, ‘dividing bacteria’ and ‘bacterial clusters’ per available pore space monitored over time, mean and standard error, n=4 chips. c) Average population density for the three classification categories after 52 days incubation of the chips *in situ* in Kenya.

To extract the distribution information, the x- and y-coordinates of the centroids of the detected single bacteria along with the estimated number and location from ‘dividing’ and ‘cluster’ was obtained from the segmented mask. The objects classified as ‘dividing’ were considered as two single bacteria allocated with random locations within 1*µ*m around the centroid of the original object. ‘Cluster’ centroids were allocated to random locations within 1*µ*mm around the original object for each estimated bacterial individual within the ‘cluster’. Subsequently, Delaunay triangulation was used to find the nearest bacteria and then calculate the distance. With this information, the index of aggregation 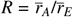 proposed by Clark and Evans in 1954 was calculated to represent the distribution of the bacteria Clark and Evans (1954). 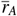 is the mean distance of each bacteria to its nearest neighbour and 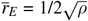 is the expected distance of even distribution where *ρ* is the density of the bacteria. The more the bacteria aggregate as clusters, the more the index of aggregation approaches zero.

## 3 RESULTS

### 3.1 Bacterial abundance

Soil chips were colonized by the bacteria deriving from the respective soil inoculum at all three locations from the arctic to the tropics. Initial examination under the microscope revealed striking differences among the bacterial communities. Here, we show a proof of concept to quantify these differences via a deep learning-based image analysis method on image data of soil chips in the laboratory or *in situ*, to obtain quantitative morphology-based bacterial population data. The number and area of the object classes ‘bacteria’, ‘dividing’, ‘cluster’ and the background class ‘microfluidic structure’ were firstly extracted to obtain the population density of each classification category over time (Figure 2 (a-b)), or as a single time point measurement (Figure 2 (c)). Chips were colonized during the first days and populations stayed at comparable levels over the duration of a month. Greenland samples were monitored longer and showed a decline over the duration of three months. In all three experiments, single ‘bacteria’ were the most commonly recognized objects whereas ‘dividing’ and ‘bacterial clusters’ were less frequent. The total bacterial density per available area in the chips differed between the communities of the three locations. Figure 3 shows cell density plotted as the mean of the total estimated number of bacteria per area for three locations over inoculation time, showing a decrease of bacterial abundance as latitude increases.

**FIGURE 3.**
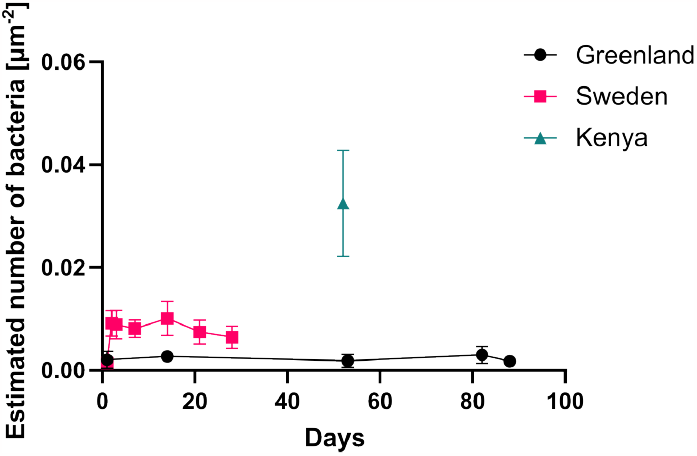
Average population density of the estimated bacterial number per area against time after inoculation in Greenland (black circle), Sweden (pink square) and Kenya (green triangle). The number of bacteria in object class ‘dividing bacteria’ is considered as two cells, and the number of bacteria in ‘bacterial clusters’ is calculated by dividing the cluster’s size by the average single bacteria size.

The algorithm allowed us to correctly capture the individual relative spatial location of each bacterial cell. We quantitatively analyzed the spatial distribution of the detected objects, where a low aggregation index number means a high level of aggregation. Figure 4 a) shows that the sample from Kenya had much more evenly distributed bacteria compared to Greenland and Sweden. Compared to the data set from the Swedish soil, a fluctuation of the index occurs over the inoculation time. Figure 4 b-d) show representative images of the spatial distributions of bacteria.

**FIGURE 4.**
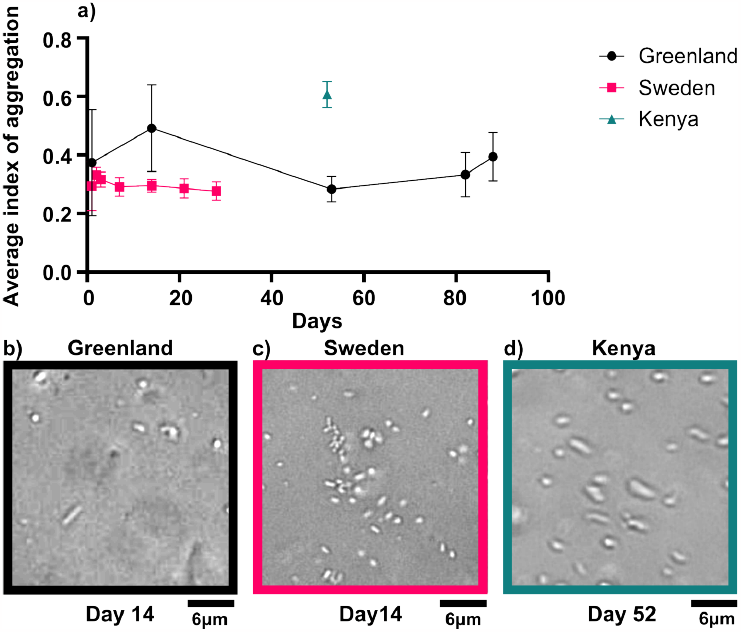
a) Aggregation index of bacterial cells, showing the deviation of the observed spatial distribution pattern from the expected random pattern. A lower index indicates more aggregated distribution. b-d) The cropped experimental images with 512*X*512 *px* for algorithm detection.

### 3.2 Morphology

The deep learning-based image analysis can instantly extract morphology information such as cross-sectional area and axis length of single object. As shown in Figure 5, a distinct discrepancy of size distribution can be found between three locations. Both the Greenland and Swedish soils have a maximum abundance of bacterial cells at 2*µm*^2^. The Swedish soil has a larger fraction of bacteria with sizes less than 4*µm*^2^, whereas the Greenland soil/biocrust has a larger subpopulation of large bacteria of 4*µm*^2^ to 6.5*µm*^2^ and around 7*µm*^2^ and generally a higher variation of cell sizes. In comparison, bacterial size in Kenya peaked at 3*µm*^2^ and the population’s size distribution is clearly shifted to the larger sizes. Mean sizes over each population are 3.32*µm*^2^, 2.50*µm*^2^, and 3.40*µm*^2^ respectively.

**FIGURE 5.**
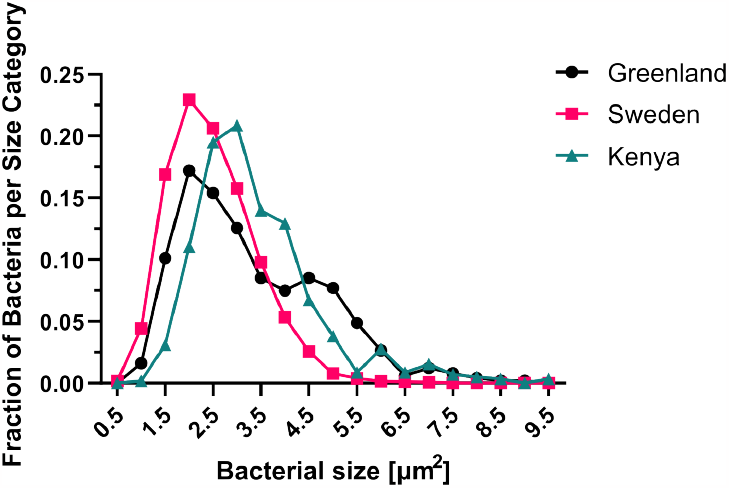
The distribution of bacterial size in Greenland (black), Sweden (pink) and Kenya (green).

To gain information about bacterial cell morphology, the ‘bacteria’ detected in the images were approximated as an ellipse, and the minor and major axis length were measured to differentiate cocci and bacilli of different shapes. As shown in Figure 6 a-c), the detected bacterial populations in three locations have distinct distribution of shapes, manifested by their distinct size along their major and minor axis. The 45^*°*^line represents the perfect circular shape. The further the bacterial major:minor axis notation is away from this line, the more elongated the bacterium becomes. The length of deviation from the axis indicates the morphological diversity of the bacterial shapes. We incorporated data from three sampling locations, which are known to exhibit discernible differences, as a validation set for our deep learning recognition model. Figure 6 revealed a significant difference in the diversity of bacteria shapes across latitudes, with the shape of bacteria being significantly less diverse in Kenya when compared to Greenland and Sweden.

**FIGURE 6.**
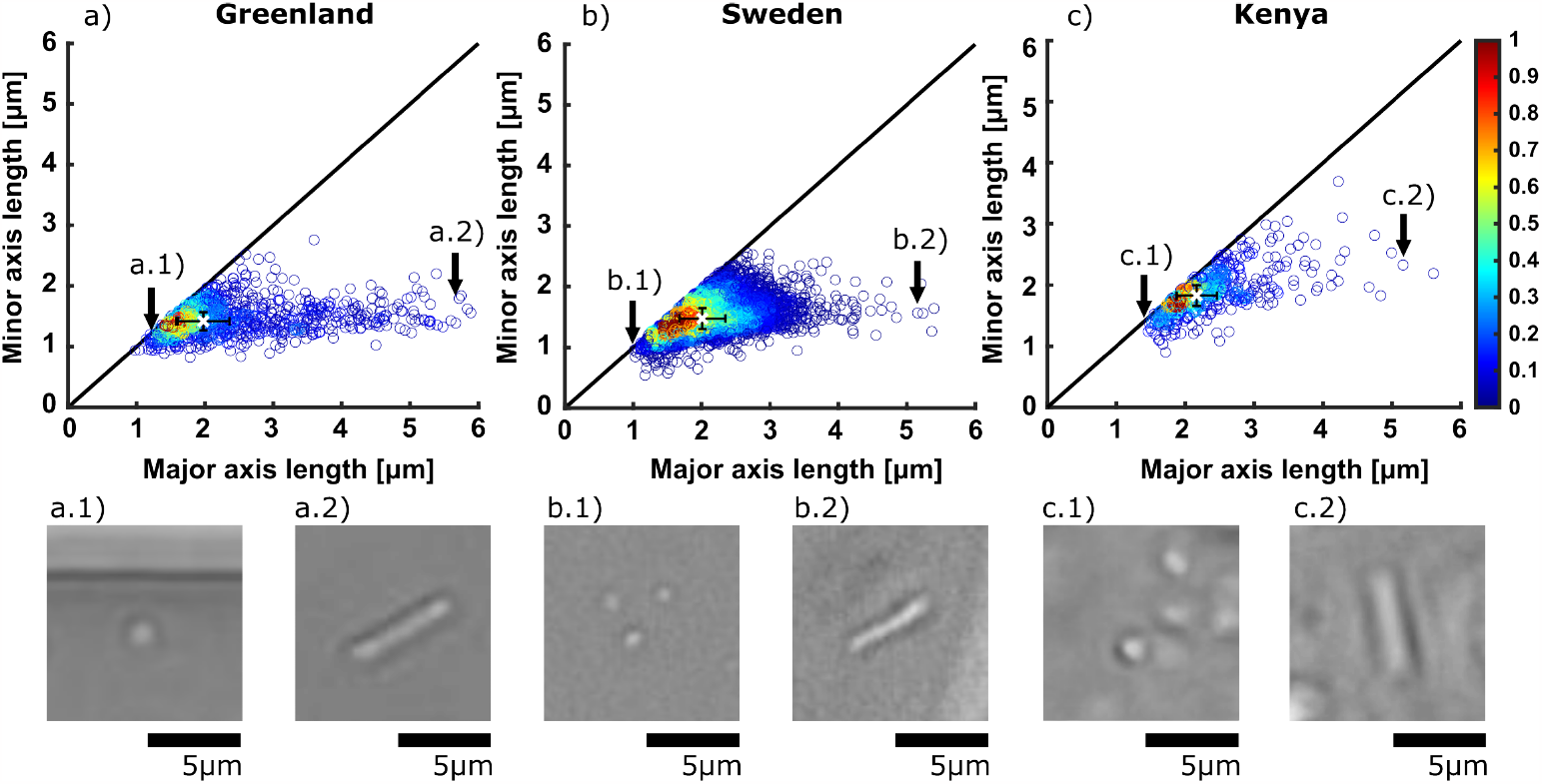
Scatterplot of minor axis length against major axis length of ellipses fitted to bacteria in chips with soils from Greenland, Sweden and Kenya. The ratio of the two axis lengths indicates how elongated or round the shape of single bacteria is. The distance to the origin point indicates the size of the bacteria in terms of the Euclidian norm of the major and minor axis. The color map on the right-hand side of the plot represent the population density of plotted points. The white cross on each plot shows the intersection of the median of the major and minor axis length, with error bars (black) of the absolute deviation of the median. a.1)-c.2) Representative example images of bacteria with corresponding shapes.

## 4 DISCUSSION

The combination of soil chips with AI-aided image analysis makes a direct investigation of soil microbial communities possible. The chips do not only allow quantification of the large microbial groups but also provide crude diversity measures within microbial groups, based on morphological phenotypes. We demonstrated this here for the characterization of the bacterial communities of three contrasting ecosystems under different experimental conditions.

We could quantify the considerable differences in bacterial abundance and community morphology variation in the different soil samples. Morphological diversity can be used as a quantitative estimate of the spatial patterns of biodiversity Roy and Foote (1997). Temperature, precipitation and soil carbon all have an effect on bacterial abundance across different spatial scales Fierer et al. (2007); Serna-Chavez et al. (2013). Fierer and Jackson suggested that a fundamental difference in biogeographical pattern exists between soil microorganisms and macroorganisms like plants and animals, and that soil microbial biodiversity is rather driven by soil pH than factors such as temperature or latitude. Bahram et al. in contrast found the highest bacterial taxonomic diversity at mid-latitudes, and falling towards the pole and the equator. Soil chips offer a global tool to morphologically examine soil microbial ecosystems, demonstrated by our analysis of samples from Greenland, Sweden, and Kenya, allowing us to study the heterogeneity of bacterial shapes that aids adaption to various environmentss Yang et al. (2016).

Apart from conventional methods of soil microbial diversity analysis such as Q-PCR, FISH, etc Dubey et al. (2020), deep learning based image analysis gives another option to directly collect data regarding bacterial abundance and morphology for work carried out on microfluidic platforms. Statistical data of distribution and morphology of live microbe in a soil-analogue environment gained from such methods is valuable for community characterization since phenotypic traits of living organisms are as important as the commonly used cell biochemical constituents. Enormous efforts have been done to promote deep learning-based methods for researchers without coding expertise von Chamier et al. (2021); Gómez-de Mariscal et al. (2021); Inés et al. (2019); Berg et al. (2019); Ouyang et al. (2019); Midtvedt et al. (2021); Sys et al. (2022). Our work showcases that the approach works under significantly different conditions, allows quantifying bacterial populations, and detects differences among bacterial communities from different locations.

Since the first attempt to measure bacterial cell sizes by Henrici et al., it has always been difficult to obtain abundance and morphology information in soil due to the methodological issues associated with visualization and recognition *in situ* Olsen and Bakken (1987). As a result, there have been limited studies on how cell abundances vary across cell size fractions and how cell morphology varies in soil Portillo et al. (2013). The abundance of individuals determines their functional role in complex communities Rivett and Bell (2018) and cell morphology is the key factor of microbe attributes as it associates with the cells’ capability of reproduction, surface attachment, motility and interaction with the environment such as nutrient intake and waste management Young (2006); Jun et al. (2018); Marshall et al. (2012); Bakken and Olsen (1987). However, the traditional manual counting including extraction, plate counting, hemocytometry and turbidimetry were proven to succumb to complexity, low precision and reliability, whereas image analysis-based methods have now been widely adopted due to their efficiency and accuracy Zhang et al. (2021); Gupta et al. (2019). The deep learning model successfully segments and classifies various objects at satisfying accuracy. AI-aided soil chip analysis does not only allow to study abundance and biodiversity measures following local treatments or large biogeographic variations, but even to investigate direct interactions of microbes with each other, or with additives to the ecosystem, such as biochar or toxins that can be added into the chips. The approach can be extended to other microbial groups including fungi, protists, invertebrates and microarthropods. The obtained cell morphology data can be useful in combination with other diversity measurements, complementing molecular taxonomic analysis with phenotypic characterization.

## 5 CONCLUSIONS

We used the well-developed deep learning algorithm Mask-RCNN to quantitatively analyse soil bacterial communities from three contrasting locations (Greenland, Sweden and Kenya), showing that the approach is capable to be used in laboratory incubations and *in situ* in field experiments. Furthermore, we demonstrated that the method is able to quantify bacterial abundance and morphology variance across latitudes. With this, we provide a proof of concept of a novel analytical technique using deep learning-based image analysis with microfluidics in order to understand and explain population and community dynamics. This approach is an important contribution to connecting controlled, high-resolution microbial analysis to field experiments and to add phenotypic analysis to soil microbial diversity studies. The concept of combining microfluidic soil chips with deep learning image detection makes it possible to obtain a more direct microbial community characterization and in the future create big databases on bacterial morphology. We demonstrated the applicability of an interdisciplinary approach combining microfluidics and AI in microbial ecology research using a widely used model, Mask R-CNN. However, it is important to note that there are other models that can be utilized for this purpose as well. With the advent of foundation models, such as large pre-trained promptable models like Segment EverythingKirillov et al. (2023), we anticipate that these models can be integrated into the whole system to enable a real time prompt-able segmentation with microbe species on the chip. The synergy of AI and microfluidics allows biologists to design and fabricate complex micro-structures, visualize the activities of migrated active microbes, and analyze characteristics of individual or communities in an automated manner. The versatility of the algorithm to adapt to different microscopy settings including a portable field microscope allows to extend studies to remote areas at a global scale.

## ACKNOWLEDGMENTS

The training and data handling were enabled by resources provided by the Swedish National Infrastructure for Computing (SNIC), partially funded by the Swedish Research Council through grant agreement no. 2018-05973. The authors wish to thank Lund University research infrastructure Correlative Image Processing and Analysis for machine learning-related consultations. The work for A.S. is partially supported by grants from eSSENCE no. 138227, Vinnova no. 2020-033375 and Rymdstyrelsen no. 2022-00282. This work is supported by a Future Research Leader grant from the Swedish Foundation for Strategic Research SSF18-0089 to E.Hammer and by BECC, the strategic research environment for Biodiversity and Ecosystem services in a Changing Climate. We acknowledge financial support by NanoLund. We acknowledge Bo Elberling and Louise Andresen Rütting for their support in sampling the Greenland biocrust, and Dries Roobroeck and Geoffrey Maritim Kimutai from the International Institute of Tropical Agriculture, Nairobi for support of the field work in Kenya.

## CONFLICT OF INTEREST

The authors declare no potential conflict of interests.

## SUPPORTING INFORMATION

16S Metabarcoding for microfluidics

## Notes

### Competing Interest Statement

The authors have declared no competing interest.

