## Supplementary material for "Bacterial community characterization by deep learning aided image analysis in soil chips"

**16S Metabarcoding: microfluidics**

**Materials and Methods**

To estimate the level of diversity of bacteria within the chips, we analyzed the amplicon sequence variant (ASV) richness of the prokaryote communities inside the chips compared to their soil inoculum. We applied metabarcoding to six samples to study bacterial and archaeal communities. Four of these samples were biological treatments, and two served as controls. The biological treatments encompassed two pairs of soil and microfluidic chips inoculated with corresponding sample soil, one from an agricultural field (GPS 55.745297° N, 13.055424°W; Micro/Soil 1) and one from a permanent meadow (GPS 55.711449°N, 13.264729°W Micro/Soil 2), respectively. The two control samples were a PCR control (PCR1 from the 16S library preparation) and a non-inoculated microfluidic chip, with the latter also acting as a control for potential DNA contamination from the DNA extraction kit. Each of these six samples underwent three technical replicates of sequencing.

*DNA extraction*

The soil chip system comprises a Polydimethylsiloxane (PDMS) slab with microstructures plasma bonded to a thin coverslip. A soil sample is inoculated on one side of the structures, and a wet tissue is used to maintain humidity. To ensure sterility, the system is enclosed in a sealed petri dish for sterilization purposes.

The sampling and DNA extraction process was conducted in a sterile hood. Initially, the sealed Petri dishes containing chips inoculated with soil were opened in the sterile hood, and 500 mg of soil sample was collected into bead tubes. The soil inoculum was removed using a spatula, and any remaining soil on the coverslip and near the chip entry was carefully wiped away using a sterilized medical cotton swab. To prevent contamination, approximately 3mm of the entry system was cut off and discarded, and the area under it cleaned. The remaining PDMS structures are meticulously sliced off with a sterilized medical scalpel to expose the colonization area. DNA sampling was performed using another sterilized cotton swab, and the swab's head was then snapped and placed into the bead tubes. The DNA extraction process followed the instruction manual of NuclseoSpin Soil (MACHEREY-NAGEL)

*Bacterial and Archaeal communities*

We amplified the V4 region of the 16S SSU rRNA gene using 515F “Parada” (5'- GTGYCAGCMGCCGCGGTAA) and 806R “Apprill” (5'-GGACTACNVGGGTWTCTAAT) primers (Apprill et al., 2015; Parada et al. 2016) and REDTaq® Master Mix (Sigma-Aldrich, Sweden). The amplification protocol was: 3 min at 94°C, followed by 40 cycles of 1 min at 95°C, 1 min at 50°C, 1.30 min at 72°C, and a final 7-min step at 72°C. We confirmed PCR products through gel electrophoresis, and the bacterial libraries underwent 2x300 bp paired-end sequencing using the MiSeq reagent kit V3 (Illumina) by Macrogen Europe (Maastricht, The Netherlands).

We processed the sequencing reads with DADA2 (version 1.29) to identify the amplicon sequence variants (ASVs) (Callahan et al., 2016). First, we removed the PCR primer sequences by trimming reads on both ends. Then, we truncated the reads to eliminate poor-quality regions (240 bp for R1 and 160 bp for R2), as indicated by DADA2 quality plots. We discarded reads containing ambiguous bases or those with expected errors exceeding two. Next, we fitted DADA2’s error models to the R1 and R2 reads to detect and rectify sequencing errors, following DADA2’s guidelines. ASVs were acquired by merging read pairs and excluding potential PCR chimeras. Using the SILVA database (release 138), which includes reference 16S rRNA gene sequences for bacteria and archaea (Quast et al., 2012), we assigned taxonomy to the ASVs.

The technical replicates of sequencing for the PCR control, the microfluidic chip control, and the soil and inoculated microfluidics were pooled. We eliminated ASVs traced to organellar genomes (specifically, 6 chloroplastic and 15 mitochondrial ASVs). Lastly, from the soil and inoculated microfluidic chips, we removed the ASVs detected in the PCR control (31 ASVs) and the non-inoculated microfluidic chips (237 ASVs).

| ASV richness | Bacteria | Archaea |
| --- | --- | --- |
| **Chip 1** | 298 | 4 |
| **Soil 1** | 753 | 10 |
| **Chip 2** | 234 | 1 |
| **Soil 2** | 878 | 15 |

**Table S1.** ASV richness of bacteria and archaea in microfluidic soil chips and respective soil inoculum.

| Micro1 vs Soil1 | Bacteria | Archaea |
| --- | --- | --- |
| **Specific micro** | 266 | 3 |
| **Shared** | 32 | 1 |
| **Specific soil** | 721 | 9 |

| Micro2 vs Soil2 | Bacteria | Archaea |
| --- | --- | --- |
| **Specific micro** | 181 | 0 |
| **Shared** | 53 | 1 |
| **Specific soil** | 825 | 14 |

**Table S2.** The number of bacterial and archaeal specific and shared ASVs in either the microfluidic soil chips or soil inoculum.
